# Discrete cortical control during quiet stance revealed by desynchronization and rebound of beta oscillations

**DOI:** 10.1101/2023.05.01.539009

**Authors:** Akihiro Nakamura, Ryota Miura, Yasuyuki Suzuki, Pietro Morasso, Taishin Nomura

## Abstract

Postural sway during quiet stance often exhibits a repetition of micro fall and the subsequent micro recovery. The classical view –that the quiet bipedal stance is stabilized by the ankle joint stiffness– has been challenged by paradoxical non-spring-like behaviors of calf muscles: gastrocnemius muscles are shortened (contract) and then stretched (relax), respectively, during the micro fall and the micro recovery. Here, we examined EEG (electroencephalogram) based brain activity during quiet stance, and identified desynchronization and synchronization of beta oscillations that were associated, respectively, with the micro fall and the micro recovery. Based on a widely accepted scenario for beta-band desynchronization during movement and post-movement rebound in the control of discrete voluntary movement, our results reveal that the beta rebound can be considered as a manifestation of stop command to punctuate the motor control for every fall-recovery cycle. Namely, cortical interventions to the automatic postural control are discrete, rather than continuous modulations. The finding is highly compatible with the intermittent control model, rather than the stiffness control model.

## 1 Introduction

The body during quiet stance is inclined slightly forward so that the mean vertical projection of the center of mass (CoM) is on average a few centimeters forward of the ankle joint [1]. This causes a tendency of the body to forward fall, due to gravitational toppling torque. For this reason, postural sway in anterior-posterior direction during quiet stance tends to exhibit a repetition of forward micro fall and the subsequent backward micro recovery. The classical view - that the quiet bipedal stance is stabilized by the ankle joint stiffness, referred to as the stiffness control [2]-has been challenged by paradoxical non-spring-like behaviors of calf muscles [3] and by neuromechanical considerations [4]: Loram et al [3] showed that gastrocnemius muscles (GA) are contracted and shortened during the micro fall, while it is relaxed and stretched during the micro recovery. More specifically, the shorting of GA during the fall is accompanied by a high-level activation (contraction) of GA, whereas the lengthening of GA during the recovery is accompanied by a low-level activation (relaxation) of GA. Notice that this process is contrary to the conventional belief of centrifugal contraction during lengthening of the calf muscles along with the micro fall. Such newly revealed fall-recovery process indicates that the recovery movement of the posture in the backward direction is not driven by active control (active contraction of calf muscles), but by inertia force *in the absence of active control*. The intermittent control, an alternative strategy to the stiffness control, asserts that upright posture is stabilized by exploiting such inertia-force-driven passive recovery dynamics that become available by switching off the active control [5–7].

In the intermittent control model, the active feedback controller, referred simply to as the neural controller, is switched on and off in a postural-state-dependent manner [6,7]. The neural controller is switched on for a period (ON period) during which the inverted-pendulum-like body falls forward (corresponding to the micro fall), where the neural controller acts as a brake to decelerate the falling motion and reverse its direction from forward to backward. When the falling motion reverses the direction, the neural controller is switched off for the subsequent period (OFF period), during which the body keeps moving backward to the upright position (to the slightly forward tilted equilibrium posture, rigorously speaking) dominantly by the inertia force with a supplemental force by the passive elasticity of the calf muscles. This is the passive recovery dynamics during the OFF period, corresponding to the micro recovery, which is the major stabilizer of the upright stance in the intermittent control model. If the timing of switching off is appropriately tuned based on the delay-affected sensory information, the body slowly approaches the equilibrium. Even in that case, however, the body eventually reverses the direction, and then falls forward, which switches on the neural controller in the next fall-recovery cycle. The intermittent control model asserts that cyclic repetition of this sequence forms a characteristic temporal pattern of postural sway. Evidence is accumulating to support the intermittent control model (e.g.,[8–10]). Particularly, we showed previously that the intermittent control model can better fit postural sway data from healthy young adults, compared to the stiffness control model, using a cutting-edge technique of Bayesian parameter inference for the model [10]. Interestingly, postural sway data from patients with Parkinson’s disease exhibiting severe postural symptoms can be better fitted by the model with less intermittency, i.e., by the model with no or a very short OFF period [10], suggesting that the appropriate placement of the OFF period is critical for the postural stabilization.

A major concern for the intermittent control model is its neural mechanisms of how the brain processes information necessary to determine the appropriate timings of the muscular activation and inactivation. Measuring cortical neural activity during postural control tasks using electroencephalogram (EEG) can approach such mechanisms. In our previous study, we examined postural responses to an impulsive backward support-surface perturbation during quiet stance and associated EEG, and identified novel brain activities [11]. Particularly, after an appearance of short latency and large amplitude synchronizations at a wide range of frequency bands in a window of a few hundred milliseconds from the perturbation [12–17], the beta-band event-related desynchronization (beta-ERD) and the subsequent long-lasting beta-band event-related synchronization (beta-ERS) occurred for a few seconds, respectively, during the forward fall response and the backward recovery response to the perturbation. The modulation of beta oscillations, i.e., the beta-ERD during movement with large muscular activations followed by the post-movement beta-ERS for the perturbed stance, is similar to those studied for the control of discrete voluntary movements of upper and lower limbs [18–21]. Thus, the beta-ERS in the late phase of the postural recovery in response to the perturbation might also be characterized as the *beta rebound* that represents the *status quo* as in the control of discrete voluntary movements, despite the difference in voluntariness and automaticity of the control. There are more differences between discrete voluntary movements and the automatic postural response to the perturbation: the discrete voluntary motor control has a clearly defined beginning and end of the movement, while the postural recovery response slowly fades out and has no clearly defined end of the movement. In this respect, beginning and end of the movement are much more unclear for postural sway during quiet stance, compared to the perturbed postural recovery response. Furthermore, there exists no specific kinematic and kinetic event, characterized by input signals, cues, nor perturbations during quiet stance. Nevertheless, if quiet stance is stabilized by the intermittent control, the motor command behind the postural sway might be punctuated by the end of each fall-recovery cycle at which the neural control is switched off.

Motivated by this hypothesis, in this study, we examine whether every fall-recovery cycle during quiet stance can be considered as a discrete motor control process in the brain. To this end, we examined whether the appearance of the OFF period (switching-off of the neural control) can be associated with a stop command for the postural control. In practice, we investigated the brain activity associated with the fall-recovery cycle, i.e., the brain activity associated with the ON and OFF of GA activation, respectively, during the micro fall and the subsequent micro recovery. If the beta-ERD and then beta-ERS were found, respectively, during the micro fall (with GA activated) and the subsequent micro recovery (with GA inactivated), it would show that the cortical motor command controlling continuous motion of the upright stance punctuate every fall-recovery cycle. In other words, cortical control during quiet stance can be considered as a discrete motor control process in the brain, rather than continuous modulations of the muscle activation.

## 2 Materials and methods

### 2.1 Participants

Twenty healthy young male participants were included in the study. None of the participants suffered from neurological disorders nor used medications that could influence posture. All participants gave written informed consent, which was approved by the ethical committee of the Graduate School of Engineering Science at Osaka University.

### 2.2 Experimental protocol

Participants were asked to stand still quietly on a force plate (BASYS, Tec Gihan Co., Ltd, Kyoto, Japan) for 150 s long. Each participant performed two trials of such quiet stance. While standing, participants folded their arm and keeping their gaze fixed on a target located 1 m in front of them at the eye level. In all trials, participants were instructed to stand upright in a relaxed state to reduce the effects of muscle-activity-derived artifacts in EEG recordings. A rest of at least 2 min was given between trials.

### 2.3 Experimental setup

Postural kinematics in the anterior-posterior (AP) and medio-lateral (ML) directions were quantified by measuring the center of pressure (CoP), for which the force plate was used to acquire time-profiles of the CoP at a sampling frequency of 1000 Hz. Only the CoP data in AP direction was used for our study. Positions of the left and the right ankle joints were measured by a three-dimensional optical motion capture system (SMART-DX, BTS Bioengineering, Milan, Italy) with a sampling frequency of 300 Hz, using light reflection markers attached on the lateral malleolus [11]. Electromyography (EMG) signals were recorded from the ankle muscles of both the left and the right legs, including the soleus, medial gastrocnemius (MG), tibialis anterior, rectus femoris and biceps femoris muscles. EMG data were recorded using wireless surface electromyograms (FreeEMG, BTS Bioengineering, Milan, Italy) with a sampling frequency of 1000 Hz. EEG signals were measured using a 64-channel mobile bio-amplifier that was placed inside a backpack and worn by the participants and a waveguard cap (eegosports, ANT Neuro, Hengelo, Netherlands). Specifically, Ag/AgCl electrodes were arranged in accordance to the International 10/10 system [22,23]. All electrodes used CPz as a reference, and AFz was used as the ground (GND). The impedance in all electrodes was controlled to be less than 25 k Ωduring the measurements [24]. All EEG signals were sampled at a sampling frequency of 2048 Hz and stored on a computer for post-processing. Data sampling of CoP, the ankle positions, EMG and EEG were started simultaneously by an identical start-trigger, where the ankle positions and EMG were sampled synchronously using a stroboscope-related clock of the motion capture system [11].

### 2.4 EEG and EMG preprocessing

Preprocessing, denoising, and analysis of EEG signals were conducted using EEGLAB v2022.0 [25]. First, the PREP pipeline was used to remove the 60 Hz line noise and rereference it to the average reference potential calculated from the less noisy electrode [26]. A zero-lag high-pass fourth-order Butterworth filter with a cutoff frequency of 1 Hz was applied to optimize signal-to-noise ratio for independent component analysis (ICA) [27]. The EEG data were downsampled to 1024 Hz to reduce the computational time required for independent component analysis. The EEG data from 10 to 150 s were used in subsequent analyses to analyze the EEG in the steady state. To remove muscle-activity and eye-blinking related artifacts from the EEG data, the ICA was performed. The artifact independent component (IC) was removed semi-automatically by the following method: Each IC was labeled using the IC label [28]. Only ICs labeled as brain activity with dipoles in the brain were used for following analysis. The removed ICs were also visually inspected.

The EMG data were band-passed from 20 to 450 Hz with a fourth-order Butterworth filter. After that, full-wave rectification was performed, and the average rectified value was calculated by applying a low-pass of 15 Hz with a fourth-order Butterworth filter.

### 2.5 Detection of micro falls and micro recoveries

Each micro fall and the subsequent micro recovery were detected by using the center of mass (CoM) time series that were estimated from the measured CoP [10,29]: First, a low-pass filter with a cutoff frequency of 6 Hz was applied to CoP using a fourth-order zero-lag Butterworth filter. Then, CoM was estimated from the filtered CoP by assuming the human body as a single inverted pendulum [29]. The CoM-positions were represented with respect to the mean position of the ankle joints in the AP direction. Namely, the mean ankle position computed from the motion captured ankle positions was set as the zero of the time-series of CoM-position in this sequel. The CoM-velocity was calculated by using a numerical differentiation (the central difference) of CoM. Positive peaks of CoM-velocity time series were detected as the instants with the fastest micro fall velocity, referred to as the micro-fall-peaks [3]. Negative peaks of the CoM-velocity time series were also detected as the instants with the fastest micro recovery velocity, referred to as the micro-recovery-peaks. Those peaks were detected from the entire CoM-velocity data of 140 s length, excluding the first 10 s to avoid transient effects. Similarly, negative and positive peaks in the CoM-position time series were also detected from the corresponding CoM-position data of 140 s length. Each negative peak of the CoM-position was considered as the beginning of a micro fall. For a given micro-fall-peak, a negative peak of the CoM-position immediately prior to the micro-fall-peak and the positive peak of the CoM-position immediately after the micro-fall-peak defined a single piece of a micro fall. This piece of micro fall was followed by a piece of micro recovery with its end determined by a negative peak of the subsequent CoM-position. A single fall-recovery cycle was defined by a set of micro fall and the subsequent micro recovery.

Several kinds of event-locked average were performed based on the set of fall-recovery cycles. For the event-locked average analysis, fall-recovery cycles associated with micro-fall-peaks that were well separated from subsequent micro-fall-peaks were utilized. More specifically, micro-fall-peaks that have no micro-fall-peaks for the subsequent 2.0 s were selected, and the fall-recovery cycles associated with such micro-fall-peaks were utilized for the event-locked average analysis. By this selection, we could collect a set of EEG activities asscoiated with pure single micro falls and the subsequent micro recoveries that were not contaminated by EEG singnals caused by the beginnings of subsequent micro falls. In the following analyses, we consider only such well-separated micro falls, micro recoveries and fall-recovery cycles.

### 2.6 Event-locked average for the fall-recovery cycles

Temporal profiles of CoM and EMG of MG as well as the time-frequency map of EEG were characterized by the event-locked average analysis. To this end, the micro-fall-peaks of CoM were selected as the triggering events. Windowed profiles ranging from -3.0 to 6.0 s (length of 9 s) with respect to the triggering micro-fall-peaks were defined as epochs, which were extracted from CoM, EMG and EEG time series data. Prior to the event-locked average, time-frequency analysis was performed for each windowed EEG time series data, for each electrode, using the wavelet transform with Morlet wavelet. The obtained time-frequency maps for each electrode were defined as the epochs of event-related spectral perturbations (ERSPs), which were used to perform the event-locked average of ERSPs [30], after a time-warp processing as described below. Since any epoch contained a full fall-recovery cycle, CoM, EMG and ERSP data only for each fall-recovery cycle were extracted, by cutting off the epoch data at both ends of the cycle. Note that length of the extracted data varied depending on the fall-recovery cycle. Then, each fall-recovery cycle data of CoM, EMG and ERSP was time-warped (i.e., either stretched or compressed along the time axis) [31,32], so that the length of the cycle became equal to the average length of the fall-recovery cycles across all cycles from all participants. Note that the time-warp was performed separately for the micro-fall-segment and the micro-recovery-segment of the cycle. That is, the average length of micro-fall-segment and that of the micro-recovery-segment across all cycles from all participants were calculated first, and then each micro-fall-segment of CoM, EMG and ERSP was time-warped, so that the length of the segment became equal to the average length of the micro-fall-segments. Similarly, each micro-recovery-segment of CoM, EMG and ERSP was time-warped, so that the length of the segment became equal to the average length of the micro-recovery-segments. Then, each time-warped fall-recovery cycle was normalized, so that the onset of micro fall corresponded to 0% of the cycle, and the end of micro recovery to 100%. Finally, the event-locked average of the normalized time-warped cycles were performed across all epochs, separately from each participant and from all participants.

Particularly, ERSP for the Cz electrode was presented as a function of time (*t*) and frequency (*f*) on the time-frequency plane. For this plot, time-average powers for EEG during each epoch was used as the baseline, and ERSP power for each frequency was represented in dB with respect to the corresponding baseline power. To examine the scalp distribution of high-beta band (20-30 Hz) modulation, the scalp distributions at the 20%, 40% and 65% of the normalized fall-recovery cycle were plotted.

To obtain the event-locked average of EMGs, time-series of EMG for each participant were normalized so that their mean values in the interval between the micro-fall-peak and the end of micro fall (the positive peak of the CoM profile) became unity. Then, the obtained EMG time-profiles were averaged across the participants.

As in the previous studies [5,6,11,33], we plotted the event-locked averaged trajectory of the fall-recovery cycle on the CoM-position vs. CoM-velocity phase plane, in which the averaged trajectory was colored according to the high-beta-band power, based on the ERSP map for the Cz electrode. Moreover, the phase portrait of the intermittent control model was computed in order to illustrate a possible association between the ON-OFF switching in the model and the high-beta-ERD/ERS appearances in the experimental EEG data. Particularly, for the phase plane of the model, the on-regions and the off-regions of the neural controller were colored by white and gray, respectively. Then, the event-locked averaged trajectory for the model-simulated fall-recovery cycles was shown, in which the trajectory was colored-coded according to the probability of the on-off selection of the neural controller. In the model simulation, we used parameter values of the model that fitted postural sway data from healthy young adults optimally in our previous study [10] using a method of the Bayesian parameter inference (the slope of the ON-OFF switching boundary was set to 0.010 [mm/s]/[mm]). See Fig. 1 and Fig. 2 of our previous study [11] for comprehensive illustrations of the intermittent control model using the phase plane analysis.

**Fig. 1.**
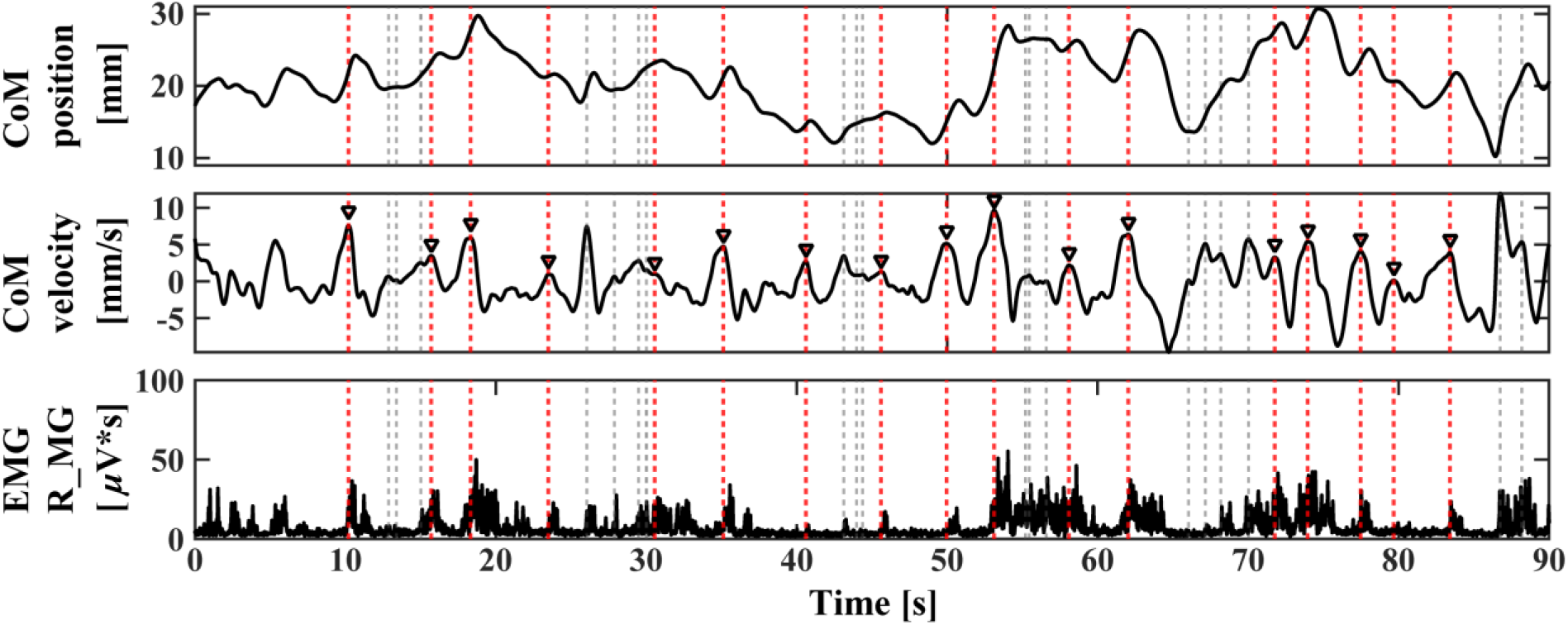
Representative postural sway data in the anterior-posterior (AP) direction during quiet stance with detected micro falls. Top and middle panels show the CoM-position and the CoM-velocity time series, respectively. The CoM-position on the top trace was represented with respect to the mean ankle joint position, i.e., the origin of the vertical axis was the mean ankle joint position. Bottom panel shows the EMG of medial gastrocnemius (MG) from the right leg. Vertical dotted lines represent the micro-fall-peaks detected as the CoM-velocity positive peaks in CoM-velocity. EEG analyses were performed only for fall-recovery cycles associated with well-separated micro-fall-peaks that are represented by the vertical red dotted lines and the open black inverted triangles.

**Fig. 2.**
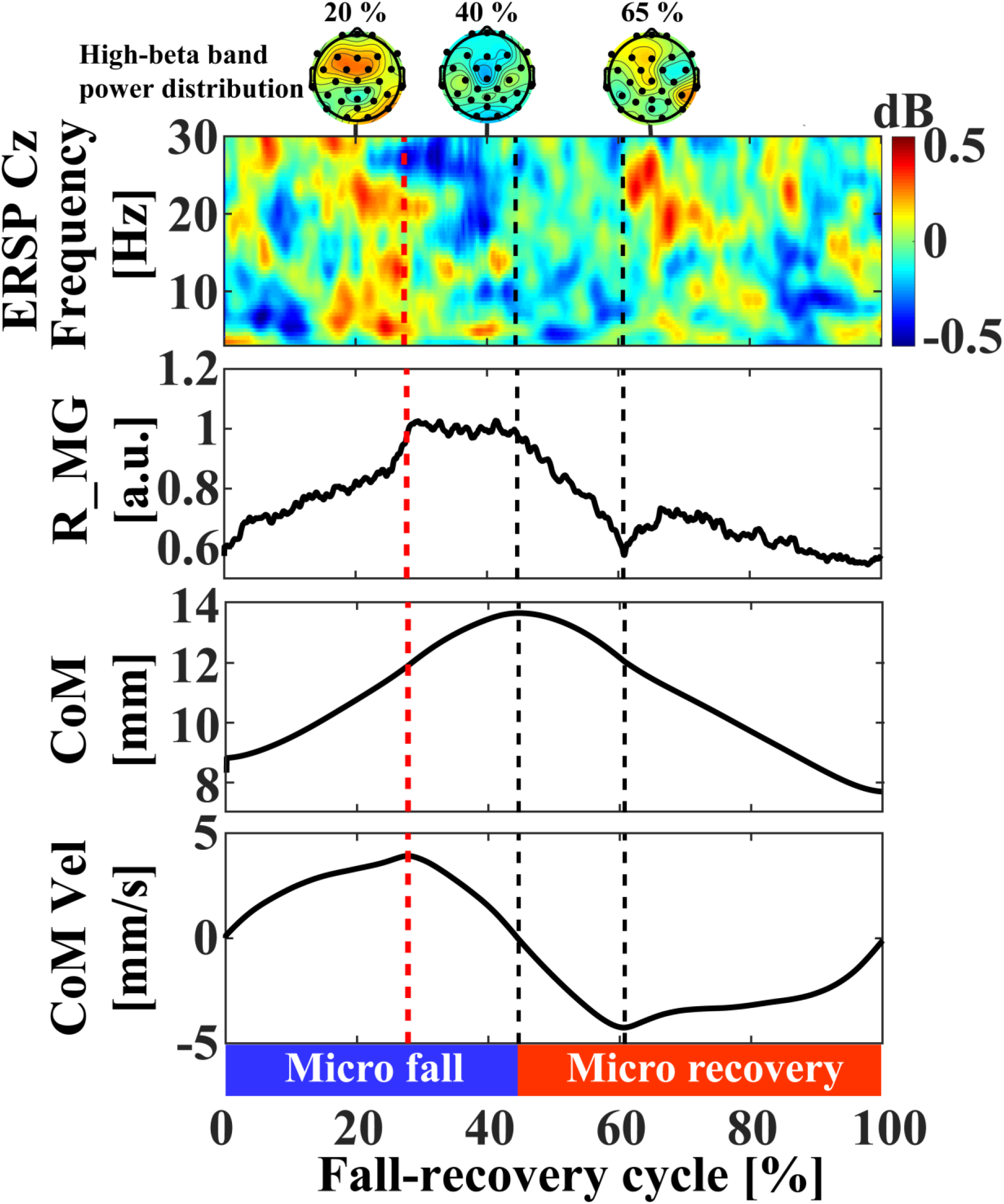
Time-warped event-locked averages of EEG-ERSP (time-frequency responses), EMG of medial gastrocnemius muscle (MG) from the right leg, the CoM-position in AP direction, and the corresponding CoM-velocity for the full fall-recovery cycle are shown in this order from the top to the bottom traces. 0% and 100% of the cycle were the beginning of the micro fall and the end of the micro recovery (the beginning of the next micro fall). The red dashed vertical line represents the micro-fall-peak defined by the positive peak of the CoM-velocity, followed by two black dashed vertical lines. The first black dashed line represents the division between the micro fall phase and the micro recovery phase. The second black dashed line represents the micro-recovery-peaks defined by the negative peak of the CoM-velocity. The CoM-position was represented with respect to the ankle joint position. Namely, the origin of the vertical axis was set to the ankle joint position in the AP direction.

To quantify the changes in the high-beta-band power within the fall-recovery cycle, the means of the high-beta band power of ERSP were calculated for three 10% windows of the fall-cycle length, which were defined by (i) immediately before the micro-fall-peak (17-27% of the fall-recovery cycle, determined after locating the micro-fall-peak at the 27% of the fall-recovery cycle), referred to as *the early fall phase*, (ii) after the micro-fall-peak (31-41% of the fall-recovery cycle, which was determined as the middle of the interval between the micro-fall-peak at 27% and the beginning of the micro-recovery at 44%), referred to as *the late fall phase*, and (iii) immediately after the micro-recovery-peak (60-70% of the fall-recovery cycle, determined after locating the micro-recovery-peak at the 60% of the fall-recovery cycle), referred to as *the late recovery phase*. Then, statistical tests were performed to compare those powers. A one-way ANOVA was conducted after verifying the normality of the data distribution using Lilliefors’ test. Subsequently, post-hoc Tukey HSD tests were performed for the three combinations at a 5% significance level [34]

## 3 Results

### 3.1 Statistics of the fall-recovery cycles

Micro falls during quiet stance occurred at the average rate of 35.1 ± 6.3 instances per minute (mean across participants). Fig. 1 illustrates micro falls detected in a representative postural sway data and the corresponding EMG time series for MG. The micro-fall-peaks that were not followed by the subsequent micro falls for 2.0 s (red dotted lines and black open inverted triangles in Fig. took place 11.4 ± 2.1 instances per minute (mean across participants). MG muscles exhibited phasic modulations that were roughly synchronous with the micro-fall-peaks.

### 3.2 Event-locked averages for the fall-recovery cycles

Fig. 2 shows the event-locked averages of EEG-ERSP, EMG of MG, CoM and CoM-velocity. In particular, the ERSP for the Cz electrode was presented together with spatio-temporal distributions of high-beta-band power on the scalp. Statistics associated with those event-locked averages were as follows: Mean length of the well-separated fall-recovery cycle was 3.61 ±1.58 s. Division ratios of the cycle for the micro-fall-segment and the micro-recovery-segment were 44% and 56%, respectively. Time profile of the CoM-position was simply bell-shaped. The EMG of MG increased during the first half of the micro fall until the micro-fall-peak (0-27% of the full fall-recovery cycle).

The ERS of EEG appeared just prior to the micro-fall-peak over a wide range of frequency band at 3-30 Hz in the ERSP for the Cz electrode. The high-beta-band component included in this ERS appeared especially in the frontal region, including the motor cortex (see the scalp map at 20% of the full fall-recovery cycle). MG activity exhibited the plateau at the maximum value in the late phase of the micro fall after the micro-fall-peak. The high-beta-ERD of EEG appeared well-coincided with the MG activation, which began slightly earlier than the micro-fall-peak. The early appearance of the high-beta-ERD split the broad-band ERS occurred just before the micro-fall-peak.

Subsequently, after the micro-fall-peak, the micro recovery began, along which MG activity decreased rapidly. For later two-thirds of the micro-recovery-segment (60-100% of the full cycle), MG activity remained at the lowest level, along which the beta rebound (the high-beta-ERS) appeared. The ERD before the micro recovery and the ERS in the late phase of micro recovery in the high-beta-band appeared both in the frontal and parietal regions including the motor cortex (see the scalp map at 40% and 65% of the full cycle).

We conducted a one-way ANOVA to compare the mean scores of beta-band power in the early fall phase, the late fall phase, and the late recovery phase. The analysis revealed a significant effect of group on the scores, F(2, 57) = 4.9, p = 0.011, MSe = 0.13. Post-hoc Tukey HSD tests indicated that beta-band power at the late fall phase (mean = -0.22, SD = 0.43) was significantly lower than that at the early fall phase (mean = -0.07, SD = 0.36) with p = 0.044. Moreover, the beta-band power at the late recovery phase (M = 0.11, SD = 0.30) was significantly higher than that at the late fall phase (p = 0.018). There was no significant difference between groups in the early fall phase and the late recovery phase (p = 0.93).

### 3.3 Event-locked averages of phase plane trajectories for the fall-recovery cycles

The event-locked average of CoM vs CoM-velocity trajectories on the phase plane for the fall-recovery cycles is shown in Fig. 3(A), exhibiting the circular curve, in which the state point moves clockwise along the curve. The blue colored high-beta-ERD started to appear at the top of the circular curve (at 12:00 o’clock), indicated by the inverted triangle that corresponds to the micro-fall-peak, and continued until the right-most portion of the curve (at 3:00 o’clock). The red colored high-beta-ERS appeared at the bottom of the curve (6:00 o’clock), indicated by the triangle that corresponds to the micro-recovery-peak. Fig. 3(B) was constructed based on numerical simulations of the intermittent control model, where the delay feedback neural controller of the model is switched on and off, respectively, in the white and the gray regions of the phase plane. The color-code between red to blue of (B) represents the probability that the neural controller is switched on: High probability was coded by blue color, and the low probability (high probability of switched-off) was coded by red color. In Fig. 3(B), the neural controller tended to be switched on from the 12:00 to 6:00 o’clock, indicated by the blue curve. At the bottom of the circular curve at 6:00 o’clock, the color of the curve was switched to red, indicating that the neural controller tended to be switched off with high probability. Overall similarity in the way of coloring the circular curves in Fig. 3(A) and (B) could be confirmed. In particular, the blue high-beta-ERD in (A) corresponded to the onset of the delayed switching-on of the neural controller, and the red high-beta-ERS corresponded to the onset of the delayed switching-off of the neural controller.

**Fig. 3.**
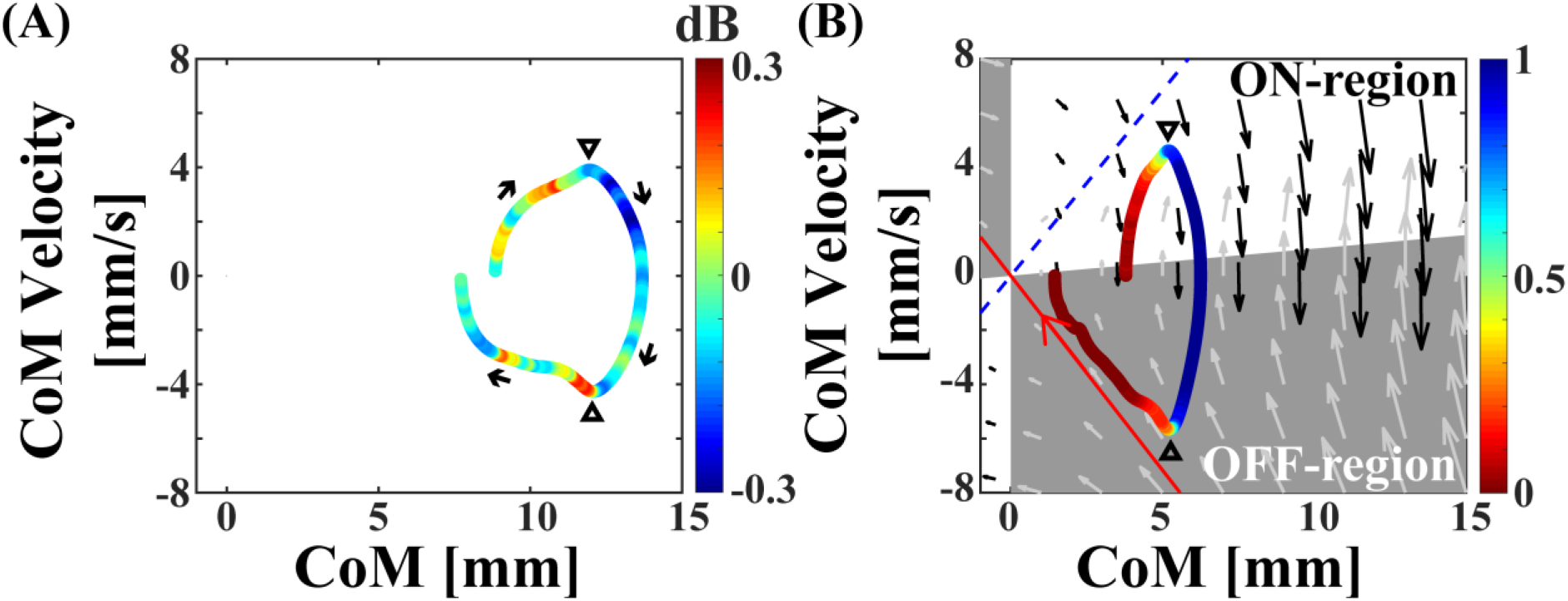
The trajectory of the time-warped event-locked average of CoM on the phase plane for the experimental data (A) and for the model-simulated data (B). The color of the trajectory indicates the average power of the high-beta-band in (A) and the probability that the neural controller is switched on in (B). In (B), the blue and the red colors represent the high and low probabilities of switched-on of the neural controller. In (A) and (B), the black inverted triangles indicate the micro-fall-peak, and the black normal triangle represent the end of the micro fall and the micro-recovery-peak. In (B), the white and the gray regions of the phase plane represent the ON-region and the OFF-region of the intermittent control model.

## 4 Discussion

In this study, we investigated the EEG-based cortical activity associated with the fall-recovery cycles (average cycle length of 3.61±1.58 s for those used for the EEG-ERSP analysis) consisting of postural sway during quiet stance. Specifically, we measured EEG activities associated with the activation (ON) and inactivation (OFF) of medial gastrocnemius muscles (MG), respectively, during the micro fall and the subsequent micro recovery. Our results showed that the high-beta-ERD appeared around the frontal and the parietal cortex in the late fall phase during which the MG exhibited the plateau at the maximum value. Then, the high-beta-ERS appeared during the late recovery phase after the MG activity decreased rapidly in the early micro recovery phase. The identification of those high-beta-ERD and high-beta-ERS was novel. Indeed, there are very few reports examining EEG in the context of postural control during quiet stance, with one exception for postural sway around the beginning of the micro recovery for postural sway in the ML direction [17], probably because degrees of synchronizations and desynchronizations associated with spontaneous postural movements during quiet stance, either phase-locked or non-locked, would be too subtle to identify them, compared to those associated with intentional [35] and/or perturbation-induced movements [11,36,37]. Although we also encountered such difficulties in this study, we have overcome the difficulties by increasing the number of fall-recovery cycles used for the event-locked average based on the time-warp methodology [31,32], while we narrowed the selection-criteria of the micro fall events for the event-locked average only to those well-separated to the subsequent micro falls in order to avoid contaminations of EEG responses associated with adjacent overlapped cycles.

Note that the switching-like activation (ON) and inactivation (OFF) of the MG, respectively, during the micro fall and the micro recovery, were reconfirmed in this study after Loram et al [3]. According to Loram et al. [3] that utilized the ultra-sound imaging of calf muscles, the MG activation during the micro fall is accompanied by the paradoxical shortening of MG, which indicates that supra-spinal mechanisms are involved in the activation and inactivation of MG, other than the mechanism of spinal stretch reflex. This finding motivated us to examine the brain activity associated with the micro fall and the subsequent micro recovery. Moreover, based on the short latency of it with respect to the micro-fall-peak, they discussed anticipatory nature of the MG activation achieved by the cerebellar internal model. In this regard, our findings of the slightly early onset of the high-beta-ERD with respect to the micro-fall-peak as well as the high-beta-ERD distributed around the frontal cortex suggest that the high-beta-ERD is associated with such predictive control.

Appearances of the beta-ERD during the micro fall with the MG activation followed by the beta-ERS during the micro recovery with MG inactivation were similar to those identified for the control of discrete voluntary movements of upper and lower limbs [18–21]. According to the widely accepted scenario for beta-band desynchronization during movement and post-movement rebound in the control of discrete voluntary movement, i.e., the beta rebound that appears after the movement as a representation of *status quo* [21,38], the beta-ERS during the micro recovery might also be characterized as the beta rebound that represents the status quo, despite the difference in voluntariness and automaticity of the control. That is, the high-beta-ERS identified in this study can be considered as a manifestation of stop command to punctuate the motor control for every fall-recovery cycle. In other words, cortical interventions to the automatic postural control, including to the mechanism of spinal reflexes, might be discrete, rather than continuous modulations. The finding is highly compatible with the intermittent control model that achieves postural stabilization by switching off the active ankle joint torque at a state-dependent appropriate timing [6,7,39], but less compatible with the stiffness control model that achieves postural stabilization using the continuous modulation of the ankle joint torque.

The broad-band ERS just before the micro-fall-peak, followed by the high-beta-ERD, then followed by the high-beta-ERS identified in this study were also quite similar to the EEG-ERSP around Cz electrode in response to an impulsive backward support-surface perturbation during quiet stance (see Fig. 3 of [11]). A noteworthy similarity among others was the identification of slightly early onset of the high-beta-ERD with respect to the micro-fall-peak, which split the broad-band ERS just before the micro-fall-peak (compare ERSP in Fig. 2 of this study and that in Fig. 3 of [11]). The splitting of the broad-band ERS caused by the high-beta-ERD could be observed much more clearly in our previous study [11] around the peak of forward fall in response to the impulsive perturbation. Note that the broad-band ERS in the perturbed response might represent the multiple spinal and cortical reflexes, including the N1 response of EEG [11–17]. Therefore, the broad-band ERS identified in this study could also represent such reflexive components in response to the micro fall even during quiet stance. Based on the similarities, including such detailed structure of the ERSP, we could say that the time-frequency pattern of EEG around Cz electrode within one fall-recovery cycle for quiet stance was a scaled-down version of that for the EEG response to the perturbation, although the response to the perturbation was a one-time transient episode that spanned about 4 seconds after the perturbation, while the fall-recovery cycle appeared repeatedly with shorter time scale and with smaller amplitudes in postural sway during quiet stance.

It was argued in our previous study [11] that the high-beta-ERS during the late phase of the postural recovery that appeared a few seconds after the perturbation might represent the active monitoring of re-afferent sensory information for the efferent motor command associated with the beta-ERD. The same argument could be applied for the high-beta-ERS in the current study. Some studies for the motor learning and control of discrete voluntary hand movements report that the beta rebound reflects a confidence level of the state estimation (state prediction) performed by internal models in the cerebellum [40]. That is, larger the beta-ERS response, the smaller the errors in the efferent-copy-based state prediction are. Thus, the beta-ERS identified in this study might also reflect an error-assessment in the postural control in comparison with the predicted posture and/or the postural verticality [41]. In the intermittent control model, it is hypothesized that the brain exploits mechanical properties of the upright body, such as passive body dynamics approaching the upright position [5–7], which requires a capability of switching off the active control at a sequence of appropriate timings. The beta-ERS identified in this study might reflect the neural activity associated with such monitoring to assess whether the control is properly switched from on to off [11]. It is likely that such information processing and associated beta-band activity are related to neural activities of the basal ganglia, as suggested by previous studies that identified attenuated beta-band desynchronization and synchronization in patients with Parkinson’s disease during upper limb motor tasks [42,43]. Moreover, the basal ganglia-cortical loop play an important role in the action selection [44]. Those reports might support an idea that the beta-band desynchronization and synchronization we found in this study reflect the information processing required for the intermittent postural control.

The neural correlates between the brain activity and the fall-recovery kinematics with the corresponding muscle activity were demonstrated in this study. However, those correlations do not necessarily represent the active monitoring and control of posture, because passive movements of ankle joints alone can modulate beta-band oscillations, including the amplitude of beta rebound [45]. Thus, our future research must go beyond the correlations to show a causality between them. One way to challenge this issue is to examine changes in the excitability of corticospinal tracts during the high-beta ERS, using, for example, the techniques of the transcranial magnetic stimulation among others [46,47].

## 5 Conclusion

We examined EEG-based brain activity during quiet stance, and identified desynchronization and synchronization of beta oscillations that were associated, respectively, with the micro fall with activation of medial gastrocnemius and the micro recovery with inactivation of medial gastrocnemius. We discussed a possibility that the beta rebound during the micro recovery is as a manifestation of a stop command to punctuate the motor control for every fall-recovery cycle, which means that cortical interventions to the automatic postural control are discrete, rather than continuous modulations.

## 7 Author Contributions

A.N. and T.N. conceived and designed the research; A.N., R.M. and Y.S. performed experiments; A.N. analyzed data; T.N. and Y.S. advised on experimental and analytical methodologies; A.N., P.M. and T.N. interpreted results; A.N. prepared figures; A.N. drafted manuscript; T.N., Y.S. and P.M. edited and revised manuscript; A.N., Y.S., R.M., P.M. and T.N. approved final version of manuscript

## References

[1] T.E. Sakanaka, M. Lakie, R.F. Reynolds, Idiosyncratic Characteristics of Postural Sway in Normal and Perturbed Standing, Front. Hum. Neurosci. 15 (2021). https://www.frontiersin.org/articles/10.3389/fnhum.2021.660470 (accessed April 10, 2023).

[2] D.A. Winter, A.E. Patla, F. Prince, M. Ishac, K. Gielo-Perczak, Stiffness Control of Balance in Quiet Standing, J. Neurophysiol. 80 (1998) 1211–1221. https://doi.org/10.1152/jn.1998.80.3.1211.

[3] I.D. Loram, C.N. Maganaris, M. Lakie, Human postural sway results from frequent, ballistic bias impulses by soleus and gastrocnemius, J. Physiol. 564 (2005) 295–311. https://doi.org/10.1113/jphysiol.2004.076307.

[4] P.G. Morasso, V. Sanguineti, Ankle Muscle Stiffness Alone Cannot Stabilize Balance During Quiet Standing, J. Neurophysiol. 88 (2002) 2157–2162. https://doi.org/10.1152/jn.2002.88.4.2157.

[5] A. Bottaro, Y. Yasutake, T. Nomura, M. Casadio, P. Morasso, Bounded stability of the quiet standing posture: An intermittent control model, Hum. Mov. Sci. 27 (2008) 473–495. https://doi.org/10.1016/j.humov.2007.11.005.

[6] Y. Asai, Y. Tasaka, K. Nomura, T. Nomura, M. Casadio, P. Morasso, A Model of Postural Control in Quiet Standing: Robust Compensation of Delay-Induced Instability Using Intermittent Activation of Feedback Control, PLOS ONE. 4 (2009) e6169. https://doi.org/10.1371/journal.pone.0006169.

[7] T. Nomura, Y. Suzuki, P.G. Morasso, A Model of the Intermittent Control Strategy for Stabilizing Human Quiet Stance, in: D. Jaeger, R. Jung (Eds.), Encycl. Comput. Neurosci., Springer, New York, NY, 2020: pp. 1–10. https://doi.org/10.1007/978-1-4614-7320-6_100698-1.

[8] M. Xiang, S. Glasauer, B.M. Seemungal, Quantitative postural models as biomarkers of balance in Parkinson’s disease, Brain. 141 (2018) 2824–2827. https://doi.org/10.1093/brain/awy250.

[9] H. Tanabe, K. Fujii, M. Kouzaki, Intermittent muscle activity in the feedback loop of postural control system during natural quiet standing, Sci. Rep. 7 (2017) 10631. https://doi.org/10.1038/s41598-017-10015-8.

[10] Y. Suzuki, A. Nakamura, M. Milosevic, K. Nomura, T. Tanahashi, T. Endo, S. Sakoda, P. Morasso, T. Nomura, Postural instability via a loss of intermittent control in elderly and patients with Parkinson’s disease: A model-based and data-driven approach, Chaos Interdiscip. J. Nonlinear Sci. 30 (2020) 113140. https://doi.org/10.1063/5.0022319.

[11] A. Nakamura, Y. Suzuki, M. Milosevic, T. Nomura, Long-Lasting Event-Related Beta Synchronizations of Electroencephalographic Activity in Response to Support-Surface Perturbations During Upright Stance: A Pilot Study Associating Beta Rebound and Active Monitoring in the Intermittent Postural Control, Front. Syst. Neurosci. 15 (2021) 48. https://doi.org/10.3389/fnsys.2021.660434.

[12] A.E. Edwards, O. Guven, M.D. Furman, Q. Arshad, A.M. Bronstein, Electroencephalographic Correlates of Continuous Postural Tasks of Increasing Difficulty, Neuroscience. 395 (2018) 35–48. https://doi.org/10.1016/j.neuroscience.2018.10.040.

[13] T. Hülsdünker, A. Mierau, C. Neeb, H. Kleinöder, H.K. Strüder, Cortical processes associated with continuous balance control as revealed by EEG spectral power, Neurosci. Lett. 592 (2015) 1–5. https://doi.org/10.1016/j.neulet.2015.02.049.

[14] T. Hülsdünker, A. Mierau, H.K. Strüder, Higher Balance Task Demands are Associated with an Increase in Individual Alpha Peak Frequency, Front. Hum. Neurosci. 9 (2016). https://www.frontiersin.org/articles/10.3389/fnhum.2015.00695 (accessed March 7, 2023).

[15] A.M. Payne, L.H. Ting, G. Hajcak, Do sensorimotor perturbations to standing balance elicit an error-related negativity?, Psychophysiology. 56 (2019) e13359. https://doi.org/10.1111/psyp.13359.

[16] J.P. Varghese, R.E. McIlroy, M. Barnett-Cowan, Perturbation-evoked potentials: Significance and application in balance control research, Neurosci. Biobehav. Rev. 83 (2017) 267–280. https://doi.org/10.1016/j.neubiorev.2017.10.022.

[17] J.P. Varghese, K.B. Beyer, L. Williams, V. Miyasike-daSilva, W.E. McIlroy, Standing still: Is there a role for the cortex?, Neurosci. Lett. 590 (2015) 18–23. https://doi.org/10.1016/j.neulet.2015.01.055.

[18] C. Neuper, G. Pfurtscheller, Evidence for distinct beta resonance frequencies in human EEG related to specific sensorimotor cortical areas, Clin. Neurophysiol. 112 (2001) 2084–2097. https://doi.org/10.1016/S1388-2457(01)00661-7.

[19] A. Stančák, B. Feige, C.H. Lücking, R. Kristeva-Feige, Oscillatory cortical activity and movement-related potentials in proximal and distal movements, Clin. Neurophysiol. 111 (2000) 636–650. https://doi.org/10.1016/S1388-2457(99)00310-7.

[20] C. Neuper, G. Pfurtscheller, Post-movement synchronization of beta rhythms in the EEG over the cortical foot area in man, Neurosci. Lett. 216 (1996) 17–20. https://doi.org/10.1016/0304-3940(96)12991-8.

[21] T. Solis-Escalante, G.R. Müller-Putz, G. Pfurtscheller, C. Neuper, Cue-induced beta rebound during withholding of overt and covert foot movement, Clin. Neurophysiol. 123 (2012) 1182–1190. https://doi.org/10.1016/j.clinph.2012.01.013.

[22] G.E. Chatrian, E. Lettich, P.L. Nelson, Ten Percent Electrode System for Topographic Studies of Spontaneous and Evoked EEG Activities, Am. J. EEG Technol. 25 (1985) 83–92. https://doi.org/10.1080/00029238.1985.11080163.

[23] V. Jurcak, D. Tsuzuki, I. Dan, 10/20, 10/10, and 10/5 systems revisited: Their validity as relative head-surface-based positioning systems, NeuroImage. 34 (2007) 1600–1611. https://doi.org/10.1016/j.neuroimage.2006.09.024.

[24] T.C. Ferree, P. Luu, G.S. Russell, D.M. Tucker, Scalp electrode impedance, infection risk, and EEG data quality, Clin. Neurophysiol. 112 (2001) 536–544. https://doi.org/10.1016/S1388-2457(00)00533-2.

[25] A. Delorme, S. Makeig, EEGLAB: an open source toolbox for analysis of single-trial EEG dynamics including independent component analysis, J. Neurosci. Methods. 134 (2004) 9–21. https://doi.org/10.1016/j.jneumeth.2003.10.009.

[26] N. Bigdely-Shamlo, T. Mullen, C. Kothe, K.-M. Su, K.A. Robbins, The PREP pipeline: standardized preprocessing for large-scale EEG analysis, Front. Neuroinformatics. 9 (2015) 16. https://doi.org/10.3389/fninf.2015.00016.

[27] I. Winkler, S. Debener, K.-R. Müller, M. Tangermann, On the influence of high-pass filtering on ICA-based artifact reduction in EEG-ERP, in: 2015 37th Annu. Int. Conf. IEEE Eng. Med. Biol. Soc. EMBC, 2015: pp. 4101–4105. https://doi.org/10.1109/EMBC.2015.7319296.

[28] L. Pion-Tonachini, K. Kreutz-Delgado, S. Makeig, ICLabel: An automated electroencephalographic independent component classifier, dataset, and website, NeuroImage. 198 (2019) 181–197. https://doi.org/10.1016/j.neuroimage.2019.05.026.

[29] P.G. Morasso, G. Spada, R. Capra, Computing the COM from the COP in postural sway movements, Hum. Mov. Sci. 18 (1999) 759–767. https://doi.org/10.1016/S0167-9457(99)00039-1.

[30] S. Makeig, Auditory event-related dynamics of the EEG spectrum and effects of exposure to tones, Electroencephalogr. Clin. Neurophysiol. 86 (1993) 283–293. https://doi.org/10.1016/0013-4694(93)90110-H.

[31] D. Li, K. Kaminishi, R. Chiba, K. Takakusaki, M. Mukaino, J. Ota, Evaluation of Postural Sway in Post-stroke Patients by Dynamic Time Warping Clustering, Front. Hum. Neurosci. 15 (2021). https://www.frontiersin.org/articles/10.3389/fnhum.2021.731677.

[32] H. Yokoyama, N. Kaneko, Y. Masugi, T. Ogawa, K. Watanabe, K. Nakazawa, Gait-phase-dependent and gait-phase-independent cortical activity across multiple regions involved in voluntary gait modifications in humans, Eur. J. Neurosci. 54 (2021) 8092–8105. https://doi.org/10.1111/ejn.14867.

[33] A. Bottaro, M. Casadio, P.G. Morasso, V. Sanguineti, Body sway during quiet standing: Is it the residual chattering of an intermittent stabilization process?, Hum. Mov. Sci. 24 (2005) 588–615. https://doi.org/10.1016/j.humov.2005.07.006.

[34] Y. Benjamini, Y. Hochberg, Controlling the False Discovery Rate: A Practical and Powerful Approach to Multiple Testing, J. R. Stat. Soc. Ser. B Methodol. 57 (1995) 289–300. https://doi.org/10.1111/j.2517-6161.1995.tb02031.x.

[35] S. Slobounov, M. Hallett, S. Stanhope, H. Shibasaki, Role of cerebral cortex in human postural control: an EEG study, Clin. Neurophysiol. 116 (2005) 315–323. https://doi.org/10.1016/j.clinph.2004.09.007.

[36] T. Solis-Escalante, J. van der Cruijsen, D. de Kam, J. van Kordelaar, V. Weerdesteyn, A.C. Schouten, Cortical dynamics during preparation and execution of reactive balance responses with distinct postural demands, NeuroImage. 188 (2019) 557–571. https://doi.org/10.1016/j.neuroimage.2018.12.045.

[37] S.M. Peterson, D.P. Ferris, Differentiation in Theta and Beta Electrocortical Activity between Visual and Physical Perturbations to Walking and Standing Balance, ENeuro. 5 (2018). https://doi.org/10.1523/ENEURO.0207-18.2018.

[38] A.K. Engel, P. Fries, Beta-band oscillations—signalling the status quo?, Curr. Opin. Neurobiol. 20 (2010) 156–165. https://doi.org/10.1016/j.conb.2010.02.015.

[39] Y. Suzuki, T. Nomura, M. Casadio, P. Morasso, Intermittent control with ankle, hip, and mixed strategies during quiet standing: A theoretical proposal based on a double inverted pendulum model, J. Theor. Biol. 310 (2012) 55–79. https://doi.org/10.1016/j.jtbi.2012.06.019.

[40] H. Tan, C. Wade, P. Brown, Post-Movement Beta Activity in Sensorimotor Cortex Indexes Confidence in the Estimations from Internal Models, J. Neurosci. 36 (2016) 1516–1528. https://doi.org/10.1523/JNEUROSCI.3204-15.2016.

[41] K. Takakusaki, Functional Neuroanatomy for Posture and Gait Control, J. Mov. Disord. 10 (2017) 1–17. https://doi.org/10.14802/jmd.16062.

[42] H.-M. Wu, F.-J. Hsiao, R.-S. Chen, D.-E. Shan, W.-Y. Hsu, M.-C. Chiang, Y.-Y. Lin, Attenuated NoGo-related beta desynchronisation and synchronisation in Parkinson’s disease revealed by magnetoencephalographic recording, Sci. Rep. 9 (2019) 7235. https://doi.org/10.1038/s41598-019-43762-x.

[43] M.C. Vinding, P. Tsitsi, H. Piitulainen, J. Waldthaler, V. Jousmäki, M. Ingvar, P. Svenningsson, D. Lundqvist, Attenuated beta rebound to proprioceptive afferent feedback in Parkinson’s disease, Sci. Rep. 9 (2019) 2604. https://doi.org/10.1038/s41598-019-39204-3.

[44] M. Ursino, F. Véronneau-Veilleux, F. Nekka, A non-linear deterministic model of action selection in the basal ganglia to simulate motor fluctuations in Parkinson’s disease, Chaos Interdiscip. J. Nonlinear Sci. 30 (2020) 083139. https://doi.org/10.1063/5.0013666.

[45] S. Walker, S. Monto, J.M. Piirainen, J. Avela, I.M. Tarkka, T.M. Parviainen, H. Piitulainen, Older Age Increases the Amplitude of Muscle Stretch-Induced Cortical Beta-Band Suppression But Does not Affect Rebound Strength, Front. Aging Neurosci. 12 (2020) 117. https://doi.org/10.3389/fnagi.2020.00117.

[46] I.A. Solopova, O.V. Kazennikov, N.B. Deniskina, Y.S. Levik, Y.P. Ivanenko, Postural instability enhances motor responses to transcranial magnetic stimulation in humans, Neurosci. Lett. 337 (2003) 25–28. https://doi.org/10.1016/S0304-3940(02)01297-1.

[47] C. Russell, N. Difford, A. Stamenkovic, P. Stapley, D. McAndrew, C. Arpel, C. MacKinnon, J. Shemmell, Postural support requirements preferentially modulate late components of the gastrocnemius response to transcranial magnetic stimulation, Exp. Brain Res. 240 (2022) 2647–2657. https://doi.org/10.1007/s00221-022-06440-5.

